# Exploring Peptide Nanodiscs Structure and Dynamics through Synergistic Approach of NMR Spectroscopy, SAS and MD Simulations

**DOI:** 10.1101/2025.09.02.673616

**Authors:** Sirine Nouri, Akseli Niemelä, Ricky Nencini, Georgios Kolypetris, Tuomas Niemi-Aro, Salla I. Virtanen, O. H. Samuli Ollila, Artturi Koivuniemi

**Author notes:** **Competing Interest Statement:** The authors declare that they have no known competing financial interests or personal relationships that could have appeared to influence the work reported in this paper.

## Abstract

Peptide nanodiscs are promising anti-atherosclerosis therapeutics, drug delivery particles and structural biology tools. However, the lack of experimental methods for structural and dynamical characterization of these particles hinders their further development. Here we integrated nuclear magnetic resonance (NMR), small-angle x-ray scattering, and small-angle neutron scattering experiments with molecular dynamics (MD) simulations to investigate the structure and dynamics of peptide nanodiscs stabilized by the apolipoprotein A-I mimetic peptide 22A with therapeutic activity against atherosclerosis. This multi-technique approach takes advantage of combining average size and shape information from small-angle scattering, peptide site-specific information from NMR spectroscopy, and interpretative power of MD simulations. Our results reveal the intrinsic polydispersity in size of peptide nanodiscs, highlighting the importance of careful interpretation when using averaged experimental parameters. Our consensus model suggests that 22A peptides are predominantly in α-helical configuration with a disordered inter-helical orientation around the lipid matrix. The terminal regions of the peptides display greater flexibility relative to the peptide core and an enhanced C-terminal exposure to solvent, which could facilitate interaction with the enzyme LCAT. Interestingly, our results indicate that peptides and lipids rotate together as a rigid body. The methodological approach described in this paper paves the way for the design of more stable and effective therapeutic nanodiscs and for the characterization of other biomolecular aggregates that are beyond the scope of current structural biology techniques.

**Significance Statement:** Nanodiscs stabilized by 22A apoA-I mimetic peptides hold significant pharmaceutical potential for treating cardiovascular diseases by mimicking HDL functions, yet their development is hindered by the difficulty of characterizing disordered biomolecular systems. Standard structural biology techniques cannot readily resolve the structure and dynamics of these peptide nanodiscs, which is essential for rational therapeutic design. Here, we integrate complementary biophysical experiments with MD simulations to establish a consensus model of 22A peptide nanodisc structure, dynamics, and interactions with biological partners at molecular resolution. Beyond advancing peptide nanodisc design, our integrative methodology provides a generalizable framework for characterizing other disordered biomolecular assemblies that are similarly challenging to conventional structural approaches.

## Introduction

Peptide nanodiscs are composed of lipid bilayer membranes stabilized by short apolipoprotein A-I (apoA-I) mimetic peptides (1, 2). Because apoA-I is the main protein component of human high- density lipoprotein (HDL) particles, these assemblies have potential therapeutic applications as HDL particle mimics with protective features against atherosclerosis (3–5). Additionally, they can be used to stabilize membrane proteins (6, 7). However, further developments of peptide nanodiscs are hindered by the lack of experimental techniques for the characterization of structural and dynamical details of such disordered biomolecular systems (8).

On the other hand, atomistic resolution structures of traditional nanodiscs – stabilized by membrane scaffold proteins (MSPs) derived from full-length apoA-I and providing a native-like environment for studying membrane proteins in solution (9, 10) – have been resolved using nuclear magnetic resonance (NMR), electron paramagnetic resonance (EPR), and transmission electron microscopy (TEM) (11), and characterized in detail by combining molecular dynamics (MD) simulations with NMR, EPR, and small angle scattering (SAS) experiments (12). However, the inherent dynamics and heterogeneity of peptide nanodiscs make them unsuitable targets for many of these standard structural biology techniques. MSP nanodiscs have a homogeneous particle size distribution, as their radius is constrained by the fixed size of the MSP belt (13), while the average size of peptide nanodiscs depends on the peptide-lipid ratio and peptide sequence (14). Furthermore, the number of molecules per peptide nanodisc cannot be directly measured from experiments although the size and molecular fractions of peptide nanodiscs would be known.

Results from solid-state NMR experiments of nanodisc forming peptides 14A and 18A on lipid bilayers suggested a double belt arrangement of these amphipathic peptides (15). On the other hand, MD simulations suggested a picket fence like organization of ELK peptides at the peptide nanodisc rim (16, 17) but suitable data for experimental validation of such model is not available. Recent results indicate that combining MD simulations with backbone ^15^N spin relaxation times from NMR, which probe the rotational dynamics of the protein backbone, can be used to construct experimentally validated atomistic-resolution dynamic models of biomolecular aggregates with heterogeneous dynamics, such as peptide micelle complexes (18) and multidomain proteins (19).

Here, we further complement this approach with small angle X-ray scattering (SAXS) and small angle neutron scattering (SANS) data to characterize dynamics and structure of nanodiscs stabilized by 22A apoA-I mimetic peptides (Esperion Therapeutics, ESP24218) (20). This peptide was optimized to activate plasma enzyme lecithin:cholesterol acyltransferase (LCAT) when bound to nanodiscs. Because LCAT esterifies cholesterol on the surface of HDL during reverse cholesterol transport and is responsible for the formation of mature HDL particles (21), 22A peptide nanodiscs bear potential therapeutic activity against atherosclerosis, which is among most common causes of death in western world (22). Detailed understanding of molecular arrangements of peptide nanodiscs provided in this work facilitates the design of more stable and effective therapeutic nanodiscs to mimic HDL functions. Furthermore, methodological advancements pave the way for characterization of other disordered biomolecular aggregates.

## Results

### Experimental characterization of peptide nanodiscs

Nanodiscs composed of 22A peptides and DMPC phospholipids with a 1:5 peptide/lipid molar ratio were prepared following our previously developed protocol (23). Dynamic light scattering (DLS) was used to characterize the size distribution profile of the particles in the sample (Fig.1A). A single peak with a volume-averaged size of 8.4 ± 1.0 nm was observed, corresponding to correctly assembled nanodiscs. No large lipid aggregates were detected. The helical content of 22A in a free form and in nanodiscs was measured by circular dichroism (CD) (Fig. 1B). In both free form and in the nanodiscs, 22A was mostly helical (79.7 ± 1.2 % and 81.7 ± 1.3 % respectively). The transmission electron microscopy (TEM) image shows the typical discoidal morphology of the nanodiscs (Fig. 1C). Size exclusion chromatography (SEC) was performed to quantify the repartition of peptide in the sample. One predominant symmetric peak was observed in the chromatogram with a retention volume around 13 mL (Fig. 1D), indicating the correct formation of nanodiscs and the absence of free peptides in solution. Based on these characterizations, we confirmed the correct formation of nanodiscs in the sample, but we also observed heterogeneity in size of the nanodiscs, as reported in other studies (13, 24). We hypothesized that it corresponds to a distribution in the number of lipids and peptides incorporated in the nanodiscs while respecting the 1:5 peptide/lipid molar ratio.

**Figure 1.**
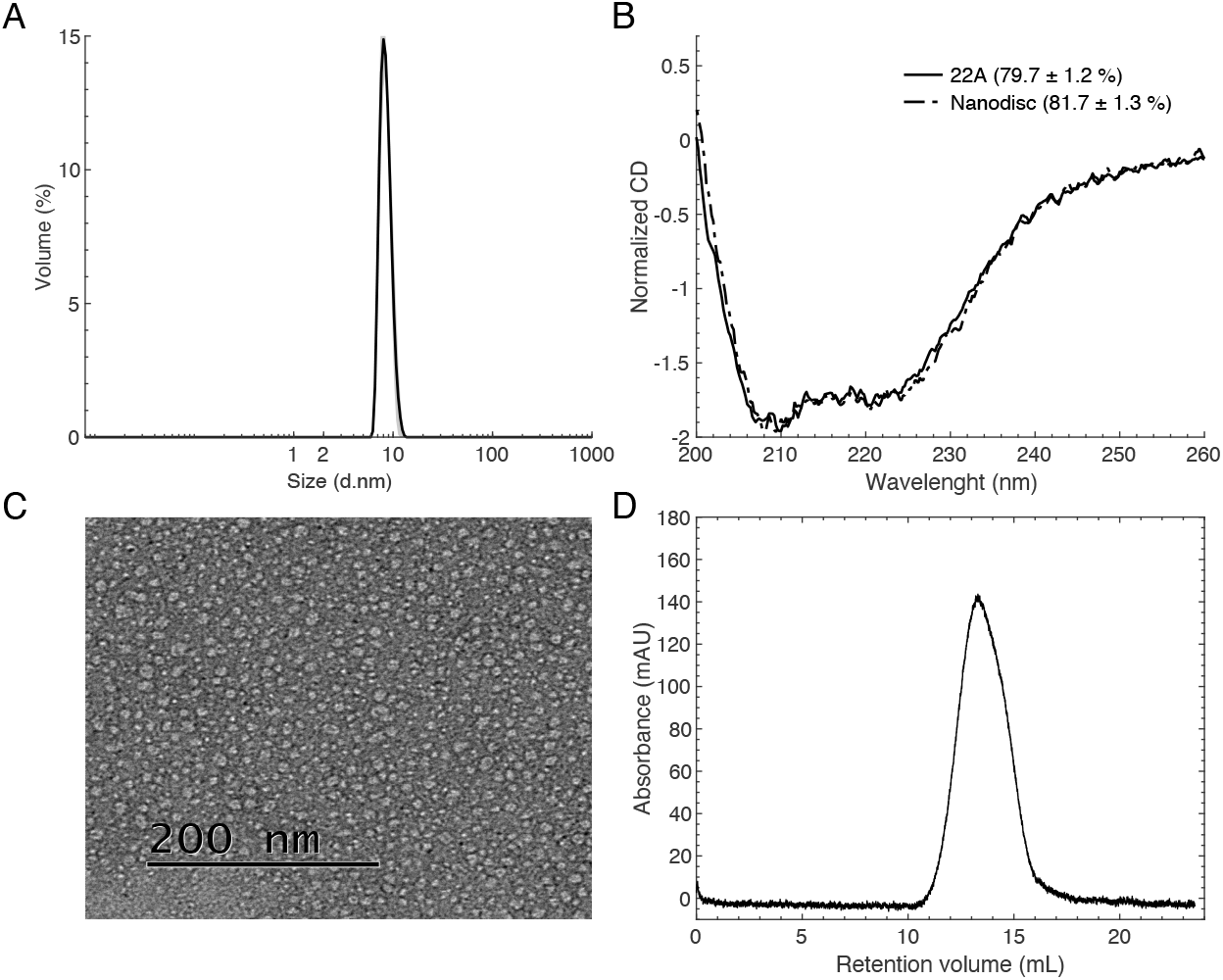
Characterization of a peptide nanodisc sample. (A) DLS measurement of the nanodiscs size. (B) Circular dichroism spectra of lipid-free 22A and nanodiscs. (C) TEM image of nanodiscs. (D) Size exclusion chromatography profile of nanodiscs.

Nanodiscs for NMR experiments were prepared using two differently labeled 22A peptides with ^15^N labels in all leucine residues [L-^15^N] or with the specifically labeled residues [R_7_-A_16_-K_18_-^15^N-^13^C]. All peaks were well resolved in ^1^H-^15^N-HSQC spectrum for both samples and each peak was successfully assigned to a residue (Fig. 2A), suggesting the absence of conformational exchange with significantly populated conformers with dynamics slower than the millisecond timescale. Nuclear spin relaxation parameters, T_1_, T_2_ and heteronuclear ^1^H-^15^N NOE, from ^15^N labeled amino acids are shown in Fig. 2B. Overall, the relaxation data are rather uniform between residues, yet showing a bell-shape dependence along the sequence, indicating a different flexibility between both termini and the internal residues.

**Figure 2.**
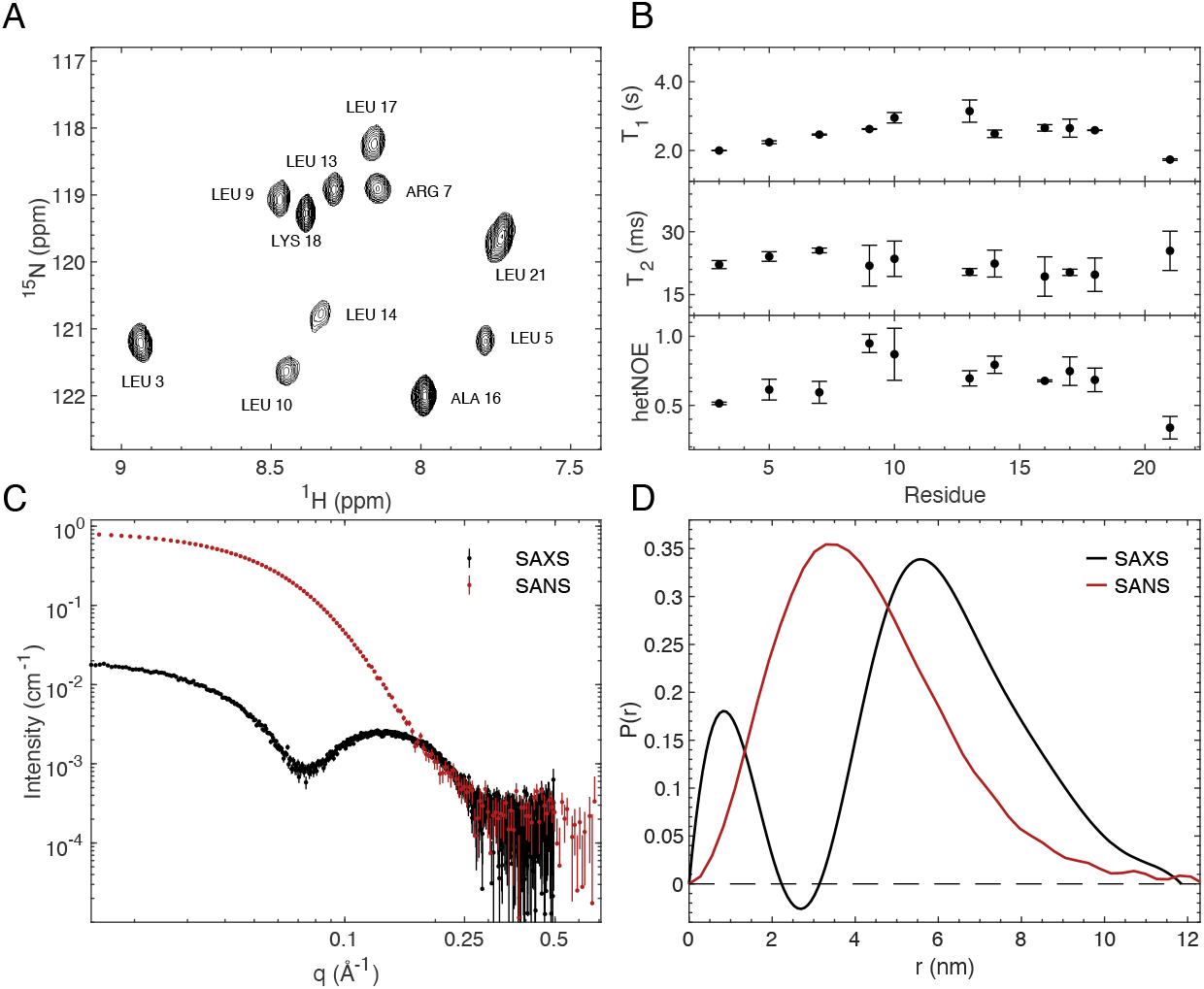
NMR and SAXS experimental data of peptide nanodiscs. (A) ^1^H-^15^N HSQC NMR spectrum. Superposition of two ^1^H-^15^N HSQC spectra from [L-^15^N] and [R_7_-A_16_-K_18_-^15^N-^13^C] labeled samples. Assignment is indicated. (B) ^15^N relaxation parameters. Data are represented as mean ± standard deviation of duplicate measurements. (C) Scattering data at 1 mg/mL peptide concentration. (D) Pair-distance distribution function derived from the scattering data. All experimental data were measured at 310 K.

SAXS data were measured for nanodisc samples at 0.12, 0.25, 0.5 and 1 mg/mL peptide concentration (Fig. S2). All scattering curves overlap at low scattering angles indicating no aggregation or interparticle effect in this concentration range. The scattering curve at 1 mg/mL peptide concentration (Fig. 2C) is used in the rest of the study as it has the lowest measurement noise. The scattering curve exhibits a flat region at low q values, a local minimum around 0.07 Å^-1^ and a broad bump at higher q values. This trend is characteristic for nanodisc solutions due to the complex scattering contrast situation, with positive excess scattering length density from the lipid headgroups and the peptides and negative excess scattering length density from the lipid hydrophobic alkyl chains (25). This is also observed in the pair-distance distribution functions P(r) (Fig. 2D), which represent the distribution of intramolecular distances within the molecular envelope. Based on SAXS P(r), the maximum distance in the nanodisc is around 12 nm, which is in good agreement with the DLS data.

SANS data were recorded on a nanodisc sample at 1 mg/mL peptide concentration in a 100 % D_2_O buffer (Fig. 2C). The 100 % D_2_O SANS data contains negative excess scattering length densities from both the peptides and the lipids. The scattering curve displays a flat region at low q values and then a decay. SANS P(r) is also indicating a maximum distance of around 12 nm in the nanodisc (Fig. 2D).

### Size distribution and number of molecules in nanodiscs from simulation-experiment integration

Results from DLS and SEC indicated that all peptides and lipids are incorporated into the nanodiscs, supporting the assumption of a 1:5 peptide/lipid ratio. However, the number of molecules in each nanodisc cannot be directly derived from experiments. Therefore, we performed MD simulations with varying numbers of lipids and peptides while maintaining the 1:5 ratio, then compared simulation results to experimental data to identify the most physically reasonable nanodisc size. We simulated nanodiscs with 50, 70, 90 and 110 DMPC lipid molecules and 10, 14, 18, and 22 22A peptide molecules, respectively, with the AMBER force field. These simulations are denoted here as MD 50, MD 70, MD 90 and MD 110. Each system was simulated for at least 3 µs and repeated three times from different initial configurations, as described in Table S1.

The calculated T_1_ and T_2_ values changed systematically with nanodisc size: T_1_ increased with larger nanodiscs, whereas T_2_ decreased (Fig. 3A). No such systematic change was observed for the hetNOE values. MD 70 exhibited the best agreement with experimental spin relaxation parameters (T_1_, T_2_, and hetNOE) compared to MD 50 and MD 90 (Fig. 3A, Table 1).

**Table 1.** Agreement between MD simulations and experiments.

|  | MD 50 | MD 70 | MD 90 | MD 110 | MD CHARMM |
| --- | --- | --- | --- | --- | --- |
| $\chi^2 - T_1$ | 0.71 | 0.29 | 1.10 | N/A | 0.68 |
| $\chi^2 - T_2$ | 2.10 | 0.39 | 0.94 | N/A | 0.68 |
| $\chi^2 - \text{hetNOE}$ | 0.70 | 0.52 | 0.54 | N/A | 0.74 |
| $\chi^2 - \text{SAXS}$ | N/A | 12.14 | 3.54 | 3.91 | 4.10 |
| $\chi^2 - \text{SANS}$ | N/A | 26.98 | 4.80 | 5.35 | 5.50 |

**Figure 3.**
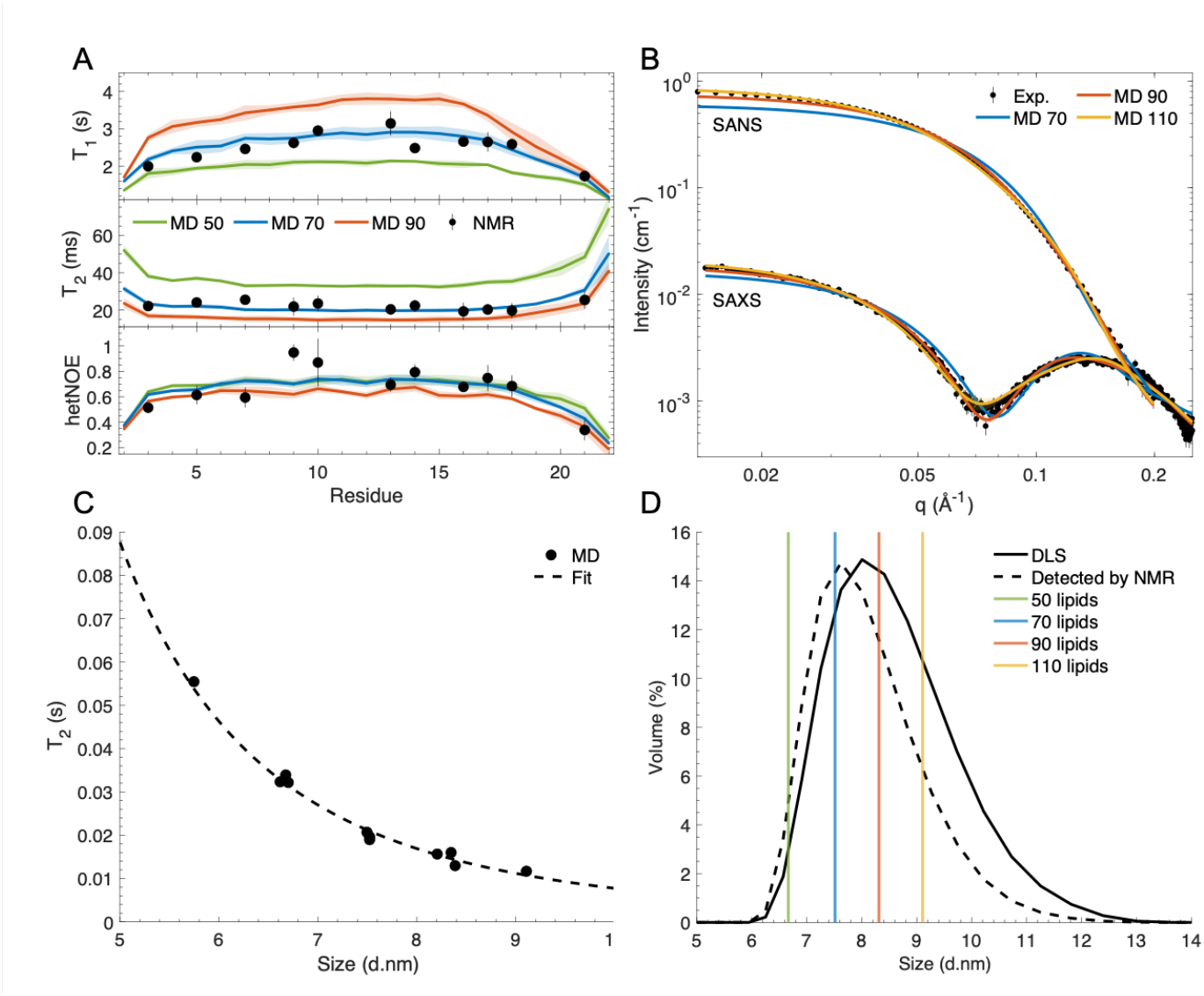
Comparing MD simulations of nanodiscs with different sizes with experiments. (A) NMR relaxation parameters. (B) SAXS and SANS curves. MD data are represented as mean ± standard deviation of triplicate simulations. (C) Correlation between T_2_ values and size of nanodiscs from MD simulations. The fitted trendline corresponds to a power function (T_2_ = 16.9 d^-3.3^, R^2^ = 0.94). (D) DLS experimental size distribution of the nanodiscs (solid black line) and rescaled distribution detected by NMR (dashed black line). Vertical lines correspond to the hydrodynamic diameters of the nanodiscs calculated from MD simulations with different number of lipids.

For SAS data, there are visible modifications of the shape of the calculated I(q) when changing the size of the nanodisc. In SAXS measurements, increasing nanodisc size produces three characteristic changes: the low-q intensity increases, the local minimum shifts to lower q values, and the broad peak at higher q values changes shape. SANS measurements show similar behavior, with larger nanodiscs exhibiting higher low-q intensities and altered curvature in the scattering decay. In contrast to NMR results, MD 90 exhibited the best agreement with experimental SAXS and SANS data in comparison with MD 70 and MD 110 (Fig. 3B, Table 1).

The discrepancy between simulation systems representing the experimental NMR and SAS data can be explained by the size polydispersity of the nanodiscs in the sample. In an NMR spectrum, each residue peak represents the averaged signal from that residue across all 22A peptides in the heterogeneous population of nanodiscs. However, the peak width of a residue is inversely proportional to T_2_ (26). Because T_2_ values decrease with increasing nanodisc size (Fig. 3A), larger nanodiscs exhibit broader NMR peaks with lower intensities than smaller ones. Therefore, smaller nanodiscs are weighted more heavily when the NMR signal is averaged over the size distribution. To quantify this effect, we first plotted the averaged T_2_ value from residues 10 to 14 against the nanodisc size (d) from MD simulations (Fig. 3C). We then fitted a power function to the data (T_2_ = 24.5 d^-3.5^, R^2^ = 0.99) and used this to rescale the DLS size distribution to that detected by NMR (Fig. 3D). This rescaling shifted the distribution maximum from the vicinity of MD 90 toward MD 70, thus explaining the observed discrepancy between nanodisc sizes determined by NMR and SAS.

In conclusion, we obtained a description of the nanodiscs from MD simulations that is consistent with structural information from NMR and SAS experiments. Our results suggest that the sample comprises nanodiscs of varying sizes, ranging from approximately 50 to over 110 lipids, with an average composition of 90 lipids and 18 22A peptides.

### Flexibility of peptides in nanodiscs

MD simulations with AMBER force field reproduce the experimentally observed bell-shape dependence of T_1_ and hetNOE along the sequence (Fig. 3A). Because this shape arises from the peptide flexibility, we analyzed peptide helicity, dynamic landscapes, per residue surface accessible solvent area (SASA) and backbone root mean square fluctuations (RMSF) from simulations for more detailed interpretation of experiments (Figs. 4, S3 and S4).

**Figure 4.**
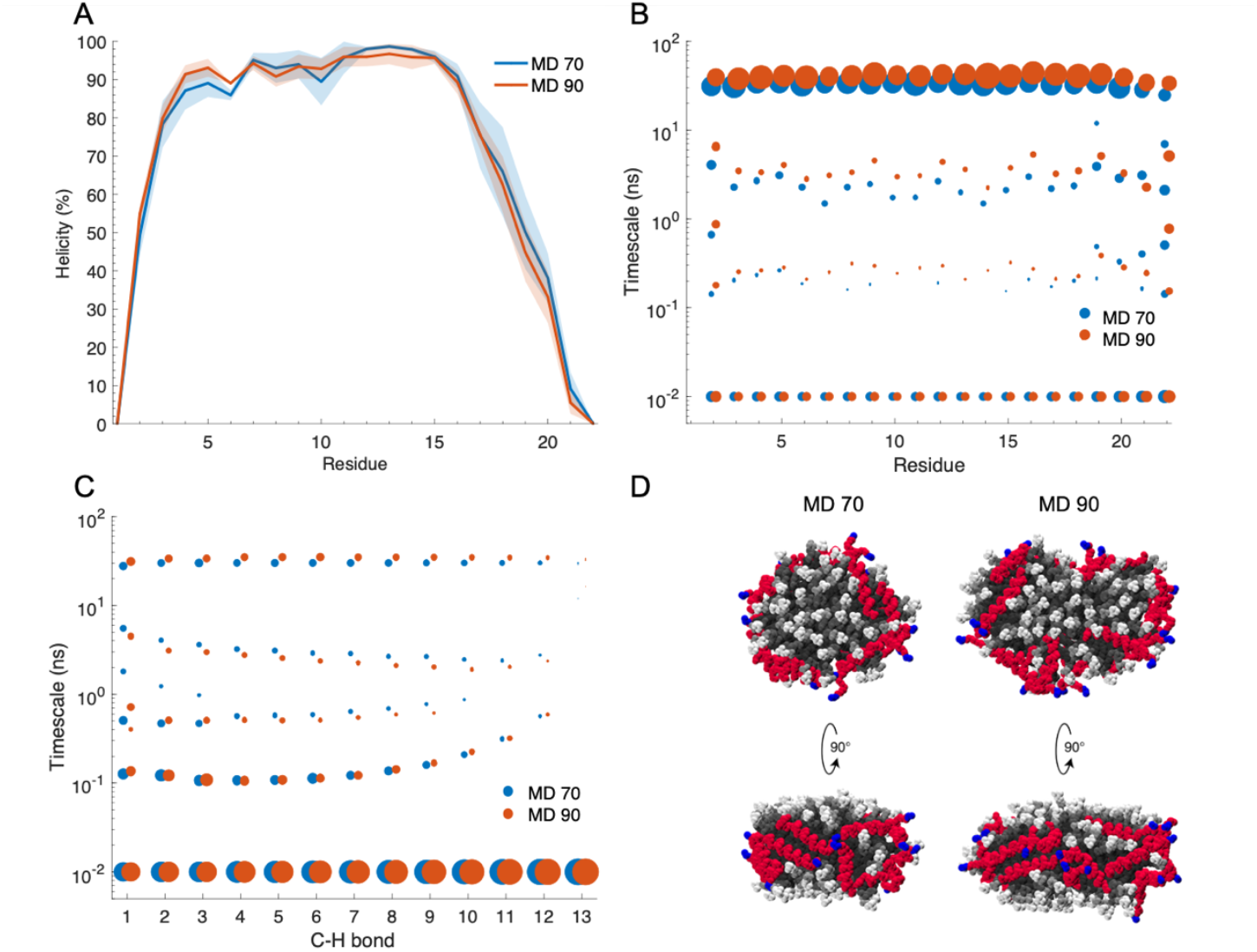
Analysis of the MD simulations. (A) Helicity per residue. MD data are represented as mean ± standard deviation of triplicate simulations. Dynamic landscapes of 22A peptides (B) and DMPC lipids (C) for one replicate of MD 70 and MD 90. C-H bond 13 refers to the terminal methyl group of DMPC phospholipids. The point sizes represent the weight of each timescale in the rotational relaxation process. (D) Snapshot of MD 70 and MD 90 simulations, top view and side view. DMPC lipids are colored in greyscale, 22A peptide in red with residue 22 in blue.

Peptides are predominantly alpha helical with low RMSF values, except N-terminal residues P1 and V2 and especially C-terminal residues K20, L21 and K22 that have lower helical propensities and higher RMSF values (Figs. 4A, S3B and S3C), suggesting pronounced flexibility in these regions. This is visible also as slightly higher weights for timescales around 100ps-5ns in the dynamic landscape profile (Figs. 4B, S4A and S4B). As expected for amphiphilic peptides, hydrophobic amino acids have lower SASA values as they face the hydrophobic acyl chains of the phospholipids (V2, L3, F6, L9, L10, L13, L14, A16 and L17), while hydrophilic amino acids are more exposed to solvent (Fig. S3A). Notably, the C-terminal residue L21 has a higher SASA value than the other leucines in the sequence and the residue K22 is highly exposed to the solvent.

For more thorough insights, we also investigated how force field parameters affect peptide flexibility by simulating the MD 70 and MD 90 systems with the CHARMM force field (denoted as MD 70 CHARMM and MD 90 CHARMM). For the NMR relaxation parameters, AMBER simulations capture the bell-shape dependence along the sequence better than CHARMM simulations, particularly for T_1_ (Fig. 5A). This can be explained by more rigid peptides predicted by CHARMM simulations with higher helical propensities than in AMBER simulations, particularly for the C-terminal end (Fig. S4C). No difference was observed for SAXS and SANS curves between AMBER and CHARMM simulations (Fig. 5B). Our results suggest that the more flexible picture predicted by AMBER is in better agreement with NMR experiments.

**Figure 5.**
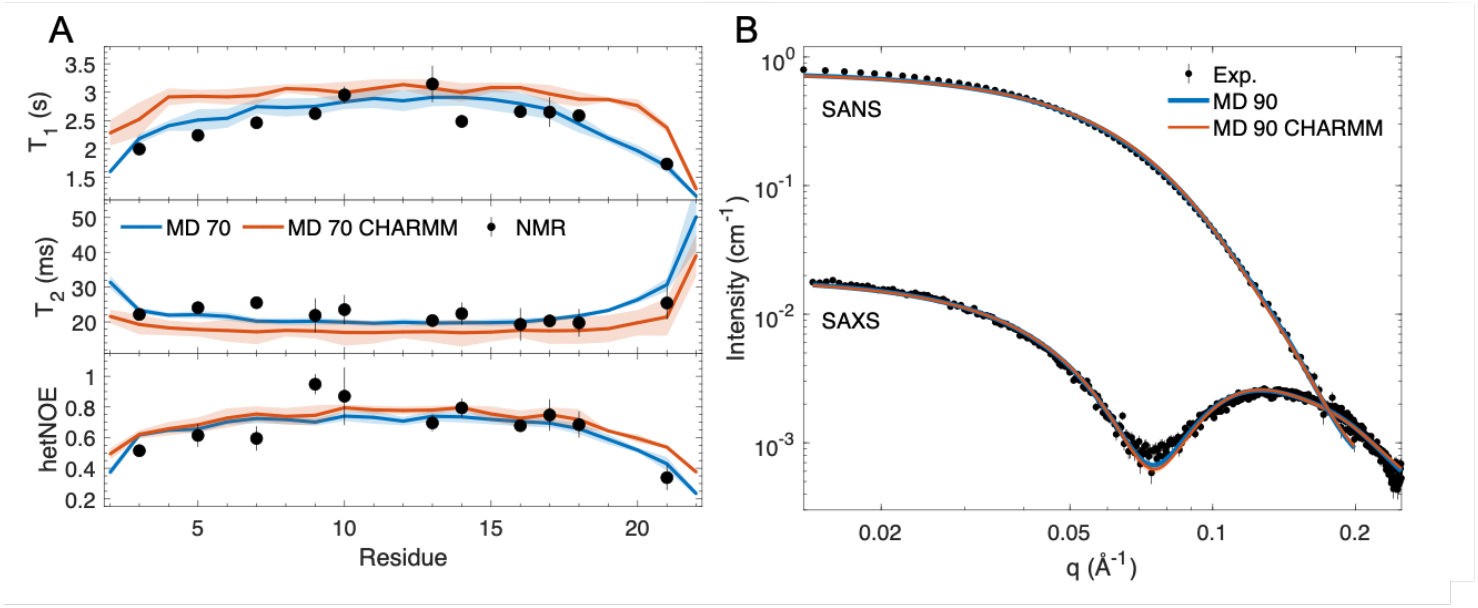
Effect of force field on the reproduction of experimental data from MD simulations. (A) NMR relaxation parameters. (B) SAXS and SANS curves. MD data are represented as mean ± standard deviation of triplicate simulations.

### Peptide orientation in nanodiscs from simulation-experiment integration

In our initial configurations, created from coarse-grained self-assembly simulations, substantial peptide ordering was not observed. Atomistic simulations retained this disorder and showed good agreement with NMR and scattering data. However, a previous simulation study suggested a picket fence-like peptide orientation, with peptides aligned parallel to the lipid bilayer normal at the nanodisc rim (17). To investigate whether the picket fence configuration would also agree with experiments, we performed MD 70 and MD 90 simulations using CHARMM and AMBER force fields with starting structures where peptides were ordered at the disc edge in a picket fence arrangement. In AMBER force field simulations, peptides did not maintain the picket fence configuration but reoriented into a more random arrangement, indicating that this conformation is unstable. Conversely, in CHARMM force field simulations, peptides remained in the picket fence configuration throughout the entire simulation.

NMR relaxation parameters were reproduced equally well by MD 70 CHARMM simulations with and without the picket fence configuration (Fig. S5A, Table S2). However, for SAXS and SANS data, the MD 90 CHARMM simulation with disordered peptides showed better agreement with experiment than the picket fence configuration (Fig. S5B, Table S2). This comparison could not be performed for AMBER simulations because the picket fence configuration was unstable.

In conclusion, AMBER simulations with more disordered peptides and unstable picket fence configurations showed better overall agreement with experimental data than CHARMM simulations, where the picket fence configuration remained stable. These results suggest that 22A peptides in nanodiscs are more likely disordered at the disc edge rather than adopting an ordered picket fence conformation. Nevertheless, we observed in Figs. 4D, S3A and S3B that the peptides arrange at the nanodisc rim with their C-termini, particularly the final residue K22, pointing toward the solvent and thus remaining accessible for interaction with LCAT. Overall, peptide behavior was similar in MD 70 and MD 90.

### Rotational dynamics of peptide nanodiscs

The rotational dynamics of peptides are characterized by a dynamic landscape plot that displays the weights of different timescales contributing to peptide rotation (Figs. 4B, S4A and S4B). These weights (α_i_ in Eq. 3) result from the fit of exponential sum to the rotational correlation functions when calculating spin relaxation times.

The rotational dynamics of 22A peptides in nanodiscs is clearly dominated by timescales around 30-40 ns having the largest weights. Such dynamic landscape is characteristic for rigid peptide structures that are mainly tumbling with the timescale corresponding to their overall rotational diffusion (27). Consistent with the peptide flexibility analysis, the C-terminal residues exhibit slightly larger weights for timescales around 100 ps–5 ns compared to other residues, indicating somewhat greater internal dynamics in this region. However, dynamic landscapes of lipid molecules are distinct from the ones of peptides as lipid C-H bond rotations are dominated by timescales faster than 100 ps (Figs. 4C, S4C and S4D). Nevertheless, most C-H bonds also exhibit a timescale of approximately 30 ns with small weight, which coincides with the overall tumbling of the 22A peptide.

In conclusion, our results suggest that lipids and 22A peptides rotate together with a common timescale of approximately 30 ns, corresponding to the overall rotational tumbling of the nanodisc. Peptides exhibit only marginal internal dynamics relative to the overall nanodisc tumbling, whereas lipids display substantial internal fluctuations, as expected for the liquid-disordered phase. This contrasts with peptide-SDS micelles, where similar analysis revealed no common timescales between peptide and SDS molecules associated with overall rotation. In that system, peptides were found to rotate in a fluid-like environment formed by the surfactants rather than as part of a rigid body (28).

## Discussion

Previous studies aiming to structurally characterize peptide nanodiscs are focused on amphipathic apoA-I mimetic 14A, 18A and ELK peptides that are similar as 22A studied here (15–17). Solid-state NMR experiments in lipid bilayers suggested double belt like arrangement for 14A and 18A peptides (15) while MD simulations suggested picket-fence like arrangement for ELK peptides (17). Here we provide more detailed picture of peptide nanodisc by integrating multiple biophysical experiments with MD simulations for DMPC nanodiscs stabilized by 22A apoA-I mimetic peptides with therapeutic functionality. Our consensus model suggests that 22A peptides are predominantly in α-helical configuration with slightly more flexible ends. With 5:1 lipid:peptide ratio, nanodiscs contain 90 lipids and 18 peptides on average, while NMR experiments see slightly smaller discs due to their faster rotational dynamics. Ordering of peptide orientations relative to one another or dimerization was not observed, and a picket fence-type arrangement appears unlikely.

On the other hand, the peptide rotational dynamics are dominated by the overall nanodisc motion, which is also observed for lipids, suggesting that all nanodisc molecules rotate together as a single unit. The internal dynamics of peptides are minimal, while lipids exist in a dynamic liquid-disordered phase. This is in contrast with peptides in surfactant micelles, where the rotation of different molecules is uncoupled (18). These results highlight the advantage of peptide nanodiscs as membrane mimics for studying membrane proteins.

Our results provide important insights on the pharmaceutical activity of 22A peptide nanodiscs. Previous MD simulations and electron microscopy experiments suggested that the C-terminal end of 22A interacts with the lid-binding groove of LCAT in the open state, thereby being relevant for the nanodisc function as LCAT activator (23). Particularly, the C-terminal lysine K22 seems to be an important residue for LCAT activation as its removal or substitution significantly reduces the cholesterol esterification activity of LCAT (23, 29). Interestingly, in our consensus model, the C-terminal end of 22A in a nanodisc is more dynamic and exposed to water than the rest of the peptide. We interpret this water exposure or more loose structure as a potentially important factor in the interaction with LCAT.

Our results pave the way for characterizing challenging biomolecular systems with heterogeneous dynamics that are difficult to study using conventional structural biology methods. Detailed experimental characterization of nanodiscs and other dynamic biomolecular complexes typically employs NMR and small-angle scattering (SAS) techniques (30, 31). NMR data provides site-specific information on dynamics and interatomic distances (32), while SAS data reveals average size and shape parameters (24). However, molecular-level interpretation of these measurements requires constructing models that are consistent with the experimental observations. Here we used MD simulations to generate atomistic resolution models that are simultaneously consistent with the dynamics detected by NMR spin relaxation experiments and structural features observed by scattering. The advantage of using MD simulations as consensus models is that they can be simultaneously validated against multiple complementary experimental techniques within a single framework. This approach is particularly useful for disordered biomolecular complexes that are beyond the scope of conventional experimental interpretation. Such targets include HDL and LDL particles, lipid nanoparticles, disordered protein aggregates, and peptide-micelle systems (18), in addition to the nanodiscs studied here.

Our results provide valuable insights into the performance of force field parameters for describing peptide nanodiscs in MD simulations. This is particularly relevant because nanodisc simulations are computationally demanding, while experimental data available for validation remains limited. MSP nanodiscs have been previously simulated at atomistic resolution with CHARMM and AMBER force fields (12, 33–36), while only CHARMM simulations are reported for peptide nanodiscs (16, 17). For 22A peptide nanodiscs, CHARMM36m produced slightly more rigid protein conformations compared to AMBER ff99SB-ILDN. The greater flexibility of AMBER ff99SB-ILDN, particularly at the termini, yielded better agreement with NMR relaxation parameters, while both force fields produced similar SAS curves, indicating comparable large-scale molecular shape and size. The picket fence-like arrangement was stable only in CHARMM36m simulations but showed poorer experimental agreement than random orientations. This stability may stem from the more rigid helical conformations characteristic of CHARMM36m. Collectively, these results suggest that AMBER simulations provide a slightly more realistic representation of 22A nanodiscs, though room remains for improvement in experimental agreement. Evaluation against additional experimental data, particularly related to lipid conformations and dynamics, would be beneficial for a more comprehensive assessment of force field performance.

In conclusion, our results combining experimental and computational data advance our understanding of the molecular-level structure and dynamics of nanodiscs formed by 22A apoA-I mimetic peptide and its interactions with LCAT enzyme. A systematic understanding of how peptide sequence variations, lipid composition, temperature, and peptide-to-lipid ratios influence nanodisc size distributions, peptide flexibility, and arrangement will facilitate the design of nanodiscs with optimal LCAT activation and therapeutic properties. This knowledge will also support precise tuning of nanodisc properties for structural biology applications. Furthermore, the developed approach shows promise for application to other disordered biomolecular aggregates, including LDL and HDL particles, lipid nanoparticles, and disordered protein condensates.

## Materials and Methods

### Sample preparation

Two apoA-I mimetic peptides 22A (PVLDLFRELLNELLEALKQKLK) with different isotopic labels in leucines [L-^15^N] or in selected amino acids [R_7_-A_16_-K_18_-^15^N-^13^C] were obtained in powder form from Peptide Protein Research Ltd (Fareham, UK). Deuterated 1,2-Dimyristoyl-sn-glycero-3- phosphocholine (DMPC) lipids were purchased in powder form from Avanti Polar Lipids Inc. (Alabaster, USA). 260 μL of a 5 mg/mL stock solution of DMPC phospholipids dissolved in chloroform was dried with a Rotavapor R-200 (BUCHI Labortechnik, Flawil, Switzerland) for one hour at room temperature. The resulting white thin lipid film was rehydrated in 1 mL of 1/50 X PBS buffer (pH 7.4). The solution was vortexed for 5 minutes and then sonicated in a water bath for one hour to obtain a clear liposome solution. 200 μL of a 5 mg/mL stock solution of labeled 22A peptides dissolved in 1/50 X PBS buffer (pH 7.4) was added. The sample was heated at 50°C for 5 minutes and then cooled to 4°C for 5 minutes. The heating-cooling cycle was repeated three times. The sample was concentrated on a Vivaspin 50k centrifugal tube (Sartorius) to reach a volume of 500 μL. 10% (v/v) D_2_O was added. The sample was transferred to a 5 mm NMR tube. The final peptide concentration is 0.75 mM and the stoichiometry is 5 lipids per peptide.

In total, four samples were produced: two samples were [L-^15^N] labeled and two samples were [R_7_- A_16_-K_18_-^15^N-^13^C] labeled. One of the [L-^15^N] labeled sample was fully characterized as described in the following section.

### Sample characterization

All experimental data were measured at 310 K, above the melting temperature of DMPC (297.1 K) where the lipids are expected to be in liquid phase.

Dynamic light scattering (DLS) (Zetasizer APS, Malvern Instruments, Westborough, MA) was used to characterize the size of the nanodiscs at 310 K. The results are shown as volume distribution of hydrodynamic diameters. Data are represented as the average of three measurements.

The helical content of lipid-free peptide and peptide in the nanodiscs was measured with a J-815 circular dichroism spectropolarimeter (Jasco). Samples were measured in a 20 mM phosphate buffer, pH 7.4 with a final peptide concentration of 0.01 mg/mL. The sample temperature was kept at 310 K during measurement using a Peltier temperature controller. 4 mL of sample were placed in a quartz cuvette with a path length of 1 cm. Data were acquired over a wavelength range of 200– 260 nm with a data interval of 0.2 nm and a scanning speed of 100 nm/min. Four scans were averaged for each sample. The percentage of helix was estimated using the CDPro analysis software with the program CONTIN (37).

The morphology of the nanodiscs was observed via transmission electron microscopy (TEM) on a Hitachi HT7800 electron microscope. The sample was diluted in a 20 mM HEPES, 120 mM NaCl, 1 mM EDTA buffer to obtain a final peptide concentration of 0.10 mg/mL. 3 μL of the sample was applied on an EM grid (200 mesh Cu + continuous carbon) for 45 seconds. The grid was washed three times with ultrapure water and blotting with filter paper before each wash. Negative staining with 2% uranyl acetate was performed first for 5 seconds and then for 60 seconds. Filter paper was used to blot the grid before the stain was applied and after the final staining. The grid was air-dried for 5 minutes. The microscope was operated at 100 kV voltage at a magnification of 40kX. The resulting pixel size was 2.8 Å/pixel.

Size exclusion chromatography was used to assess the purity of the nanodiscs. The sample was eluted through a Superdex 200 Increase column by 1X PBS (pH 7.4) with an isocratic flow rate of 0.4 mL/min and detected at 200 nm.

### NMR spectroscopy

NMR measurements were performed on a Bruker 850 MHz Avance III HD spectrometer, equipped with a cryogenically cooled TCI triple-resonance probe, at the Finnish Biological NMR Center. All experiments were measured at 310 K. [^15^N,^1^H]-HSQC, [^1^H,^1^H]-TOCSY, and ^1^H NOESY experiments were used for the assignment of the labeled peptide residues, as described previously (18). The ^1^H NOESY mixing time was set to 280 ms. TROSY-based NMR pulse sequences (38) were used to measure the longitudinal (T_1_) and transverse (T_2_) relaxation times and the heteronuclear nuclear Overhauser effects (NOE) of ^15^N nuclei in N-H groups. Series of two- dimensional spectra were recorded with different delays to determine relaxation rate constants. The following delays were used: T_1_ delays of 50, 150, 300, 500, 800, 1000, 1500 and 3000 ms and T_2_ delays of 16, 32 and 48 ms. Two heteronuclear NOE spectra were recorded with and without proton saturation during relaxation delay. All data sets were acquired using 128 scans and 32 increments with a recycle delay of 3.5 s. T_1_, T_2_ and hetNOE data were measured from all four samples providing two independent values for each spin relaxation observable.

Spectra were processed using nmr-Pipe (39). CcpNmr (40) was used to determine the intensities, in arbitrary units, for the amide ^1^H-^15^N cross peaks by measuring the height of the peaks. Intensities of cross peaks were fitted to single exponential functions to determine relaxation rate constants T_1_ and T_2_ (Fig. S1). Heteronuclear NOE values were obtained by taking the ratio of the intensities measured in the spectra collected with and without proton saturation. Error bars were calculated as standard deviation from relaxation measurements done on two independently prepared samples.

### SAXS and SANS measurements

SAXS and SANS data were performed at the BM29 BioSAXS beamline (41) at the European Synchrotron Radiation Facility (ESRF) and at the D22 beamline at the Institute Laue-Langevin (ILL), respectively, in Grenoble, France. The scattering intensity, I(q), was measured as a function of q = 4πsin(θ)/λ, where q is the scattering vector, 2θ is the scattering angle, and λ is the X- ray/neutron wavelength. Both the SAXS and SANS measurements were done at 310 K.

For the SAXS measurements, the nanodisc sample was measured in 1/50 X PBS buffer (pH 7.4), at four peptide concentrations (0.12, 0.25, 0.5 and 1 mg/mL) to avoid inter-particle interactions, at an X-ray energy of 12.5 keV. With λ = 0.99 Å and a sample-detector distance of 2.8 m, data were obtained in the q-range from 0.0085 Å^-1^ to 0.494 Å^-1^. Ten SAXS frames of 1 second were averaged, background subtracted and normalized to absolute scale units (cm^-1^) using water as calibration standard (42) with the data reduction software ScÅtter (43).

For the SANS measurements, the usual H_2_O based buffer was substituted with a corresponding buffer with 100% D_2_O. The nanodiscs sample at 1 mg/mL peptide concentration was placed in a Hellma quartz cuvette (1 mm path). The instrument was set up with a neutron wavelength of 6 Å +/- 10%, a collimation length of 5.6 m and two different sample-detector distances, 1.4 m and 5.6 m, yielding data in the q-range from 0.0094 Å^−1^ to 0.69 Å^−1^. The data reduction was performed according to standard ILL procedures by applying corrections for detector sensitivity, electronic noise, and empty cell scattering. The intensity was normalized to absolute scale units (cm^-1^) using the flux of the direct beam. Buffer subtraction was done with IGOR Pro.

Pair-distance distributions were calculating using GNOM from the ATSAS package (44).

### MD simulation

To create initial structures for atomistic simulations, we first initiated coarse-grained Martini 3 simulations (45) with 30, 50, 70, 90 and 110 DMPC lipid molecules and 6, 10, 14, 18, and 22 22A peptide molecules, respectively, maintaining the 1:5 peptide:lipid ratio corresponding to experiments. These systems were simulated for 10 μs. After placing the molecules into a random configuration for each replicate, the molecules spontaneously self-aggregated into nanodisc particles within 1 μs with DMPC lipid bilayer in the core and 22A peptides on the edges. The final snapshots were converted to atomistic resolution with CHARMM-GUI (46). AMBER systems used Amber ff99SB-ILDN and Lipid17 force fields (47, 48), while CHARMM systems used CHARMM36m for proteins (49) and CHARMM36 for lipids (50). TIP4P-Ew water model (51) was used in all systems because it predicts correct viscosity and thereby enables reproduction of experimental spin relaxation data (27). Atomistic simulations were performed using Gromacs version 2020.5 (52). A timestep of 2 fs was used, the temperature was coupled using v-rescale thermostat to 310 K (53), the pressure was set to 1 bar using isotropic Parrinello–Rahman barostat (54), PME was used to calculate electrostatic interactions at distances longer than 1.0 nm (55), and Lennard-Jones interactions were cut off at 1.0 nm.

To create initial configurations for the simulations with picket fence configurations, DMPC lipid bilayers of 20 x 20 nm were created and equilibrated using CHARMM-GUI. Circular cross sections with 70 or 90 lipids were then taken from the equilibrated bilayer. For replicates, the cross sections were taken from different locations. 22A was placed in a circle around the sections in parallel picket fence configuration. The circle radius was 3.5 nm for 70 lipid systems and 4.0 nm for 90 lipid systems.

All systems were first equilibrated for 500 ps with 200 kJ mol^-1^nm^-2^ and 50 kJ mol^-1^nm^-2^ restraints on protein backbone and side chains respectively. Then only protein backbone was restrained with 50 kJ mol^-1^nm^-2^ for 500 ps, and finally without restraints the systems were run for 100 ns. The production systems which were analyzed were run for at least 1 µs.

### Calculating spin relaxation times from MD simulations

Spin relaxation times T_1_, T_2_ and hetNOE values were calculated from simulations as described previously (18, 27). Briefly, the spectral density, J(w), was calculated as the Fourier transformation of the second-order rotational correlation function C(t) of an N-H peptide bond:

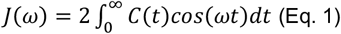

C(t) was calculated using the Gromacs gmx rotacf module from trajectories with snapshots generated every 10 fs as

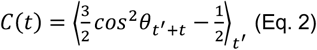

where θ is the angle between the bond in time t′ and time t′ + t. For the Fourier transformation and dynamic landscape analysis, the correlation function C(t) was fitted with a large set of N = 300 exponential functions with predefined timescales τ_i_ logarithmically spaced between 10 fs and 1 μs as

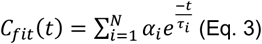

where α_i_ is the weight of a fitted pre-defined timescale τ_*i*_. The same methodology was used to calculate the weights for C-H bonds of DMPC phospholipids.

A simulation of a single molecule produces rotational correlation functions with reliable statistics up to approximately one-hundredth of the total simulation time (56), which corresponds to 10 ns for our 1 μs simulation. However, our simulations exhibit rotational timescales extending to 30-40 ns. Our simulations contain multiple peptides over which correlation functions are averaged, thereby improving statistical quality and enabling reliable analysis beyond 10 ns. When correlation functions were extended beyond the point where statistical fluctuations became significant, exponential fitting produced spurious components with artificial timescales of 1 μs. We therefore determined the maximum analysis time for each system (either 50 ns or 100 ns, as specified in Table S1) such that these artificial timescales did not influence our conclusions. The spurious long-timescale components were excluded from the calculation of spin relaxation times.

The analytical solution of the Fourier transform was used to calculate the spectral density as

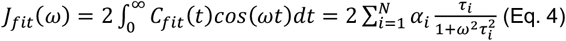

Spectral density was then substituted to Redfield equations (57, 58) to calculate spin relaxation times and hetNOE values at the magnetic field of 850MHz corresponding the experimental data. Average spin relaxation times over three replicates are reported and error bars are calculated as the standard deviation.

### Calculating SAXS and SANS from MD simulations

Theoretical SAXS and SANS profiles were calculated from MD simulations using CRYSOL (59) and CRYSON (60), respectively. PDB structures were extracted every 10 ns from the simulations and used to calculate scattering curves for the q-range from 0.0085 Å^-1^ to 0.25 Å^-1^ for SAXS and from 0.0094 Å^-1^ to 0.20 Å^-1^ for SANS. Wide-angle regime was not considered as the noise was significant in the experimental curve. Three parameters (the average displaced solvent volume per atomic group, the contrast of the hydration shell and the relative background) were fitted to the experimental curve to account for the solvation layer around the nanodisc and the buffer subtraction. The maximum order of harmonics was set to 30, the electron density of the solvent to 0.334 e/Å^3^ and the order of Fibonacci grid to 18. Hydrogen atoms were explicitly included.

The quality of the fit to experimental data is determined by the chi-squared: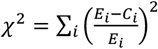, with E_i_ the experimental values and C_i_ the calculated values from MD simulation.

### Calculating helicity, SASA, RMSF and hydrodynamic diameter from MD simulations

Per residue helicities were calculated with the DSSP module of MDAnalysis (61, 62). Solvent accessible surface areas (SASA) per residue for all peptide atoms were calculated with Gromacs gmx sasa module using standard settings. Root mean square fluctuations (RMSF) were calculated with Gromacs rmsf module by fitting the peptide backbone atoms of each peptide individually to an ideal alpha helical backbone model. Individual per residue RMSF profiles were averaged per system. These analyses were performed to trajectories with snapshots generated every 1 ns. The average and standard deviation between replicate systems are shown for helicity, SASA and RMSF plots. The hydrodynamic diameter was calculated using Hydropro10 software (63) for snapshots generated every 100 ns.

## Acknowledgments

We acknowledge the European Synchrotron Radiation Facility (ESRF) for provision of synchrotron radiation facilities under proposal number MX-2582 and we would like to thank Mark Tully for assistance and support in using beamline BM29. We acknowledge the Institut Laue-Langevin (ILL) for provision of neutron radiation facilities and we would like to thank Anne Martel for assistance and support in using beamline D22. The facilities and expertise of the HiLIFE NMR unit at the University of Helsinki, a member of Instruct-ERIC Centre Finland, FINStruct, and Biocenter Finland are gratefully acknowledged. We acknowledge CSC – IT Center for Science for computational resources. This work was supported by The Academy of Finland [grant numbers 315596, 350636, 353815, 356568].

## Notes

### Competing Interest Statement

The authors have declared no competing interest.

### Summary of Updates

The abstract, introduction, result and discussion sections have been updated for clarification. Supplemental data have been added on the dynamic landscape of the nanodiscs.

## References

1. D. O. Osei-Hwedieh, M. Amar, D. Sviridov, A. T. Remaley, Apolipoprotein mimetic peptides: Mechanisms of action as anti-atherogenic agents. Pharmacology & Therapeutics 130, 83–91 (2011).

2. M. Navab, et al., Peptide mimetics of apolipoproteins improve HDL function. Journal of Clinical Lipidology 1, 142–147 (2007).

3. B. A. Kingwell, M. J. Chapman, A. Kontush, N. E. Miller, HDL-targeted therapies: progress, failures and future. Nat Rev Drug Discov 13, 445–464 (2014).

4. G. M. Anantharamaiah, D. Goldberg, Eds., Apolipoprotein Mimetics in the Management of Human Disease (Springer International Publishing, 2015).

5. D. Sviridov, A. T. Remaley, High-density lipoprotein mimetics: promises and challenges. Biochemical Journal 472, 249–259 (2015).

6. S. R. Midtgaard, et al., Self-assembling peptides form nanodiscs that stabilize membrane proteins. Soft Matter 10, 738–752 (2014).

7. M. Zhang, et al., Reconstitution of the Cyt b5 –CytP450 Complex in Nanodiscs for Structural Studies using NMR Spectroscopy. Angew Chem Int Ed 55, 4497–4499 (2016).

8. A. O. Elzoghby, et al., Nanodiscs: Game changer nano-therapeutics and structural biology tools. Nano Today 53, 102026 (2023).

9. F. Hagn, M. L. Nasr, G. Wagner, Assembly of phospholipid nanodiscs of controlled size for structural studies of membrane proteins by NMR. Nat Protoc 13, 79–98 (2018).

10. U. Günsel, F. Hagn, Lipid Nanodiscs for High-Resolution NMR Studies of Membrane Proteins. Chem. Rev. 122, 9395–9421 (2022).

11. S. Bibow, et al., Solution structure of discoidal high-density lipoprotein particles with a shortened apolipoprotein A-I. Nat Struct Mol Biol 24, 187–193 (2017).

12. T. Bengtsen, et al., Structure and dynamics of a nanodisc by integrating NMR, SAXS and SANS experiments with molecular dynamics simulations. eLife 9, e56518 (2020).

13. N. Skar-Gislinge, N. T. Johansen, R. Høiberg-Nielsen, L. Arleth, Comprehensive Study of the Self-Assembly of Phospholipid Nanodiscs: What Determines Their Shape and Stoichiometry? Langmuir 34, 12569–12582 (2018).

14. D. Xu, et al., Reconfigurable Peptide Analogs of Apolipoprotein A-I Reveal Tunable Features of Nanodisc Assembly. Langmuir 39, 1262–1276 (2023).

15. E. S. Salnikov, G. M. Anantharamaiah, B. Bechinger, Supramolecular Organization of Apolipoprotein-A-I-Derived Peptides within Disc-like Arrangements. Biophysical Journal 115, 467–477 (2018).

16. M. Pourmousa, R. W. Pastor, Molecular dynamics simulations of lipid nanodiscs. Biochimica et Biophysica Acta (BBA) - Biomembranes 1860, 2094–2107 (2018).

17. R. M. Islam, et al., Structural properties of apolipoprotein A-I mimetic peptides that promote ABCA1-dependent cholesterol efflux. Sci Rep 8, 2956 (2018).

18. R. Nencini, et al., Probing the dynamic landscape of peptides in molecular assemblies by synergized NMR experiments and MD simulations. Commun Chem 7, 28 (2024).

19. A. E. Sandelin, et al., Quality Evaluation Based Simulation Selection (QEBSS) for analysis of conformational ensembles and dynamics of multidomain proteins. Commun Chem 8, 241 (2025).

20. J.-L. Dasseux, et al., Apolipoprotein A-I agonist and their use to treat dyslipidemic disorders. (1999).

21. D. Li, S. Gordon, A. Schwendeman, A. T. Remaley, “Apolipoprotein Mimetic Peptides for Stimulating Cholesterol Efflux” in Apolipoprotein Mimetics in the Management of Human Disease, G. M. Anantharamaiah, D. Goldberg, Eds. (Springer International Publishing, 2015), pp. 29–42.

22. W. Yuan, et al., Systematic evaluation of the effect of different apolipoprotein A-I mimetic peptides on the performance of synthetic high-density lipoproteins in vitro and in vivo. Nanomedicine: Nanotechnology, Biology and Medicine 48, 102646 (2023).

23. L. Giorgi, et al., Mechanistic Insights into the Activation of Lecithin–Cholesterol Acyltransferase in Therapeutic Nanodiscs Composed of Apolipoprotein A-I Mimetic Peptides and Phospholipids. Mol. Pharmaceutics 19, 4135–4148 (2022).

24. V. Graziano, L. Miller, L. Yang, Interpretation of solution scattering data from lipid nanodiscs. J Appl Crystallogr 51, 157–166 (2018).

25. N. Skar-Gislinge, et al., Elliptical Structure of Phospholipid Bilayer Nanodiscs Encapsulated by Scaffold Proteins: Casting the Roles of the Lipids and the Protein. J. Am. Chem. Soc. 132, 13713–13722 (2010).

26. J. Cavanagh, W. J. Fairbrother, A. G. Palmer III, M. Rance, N. J. Skelton, Protein NMR Spectrocopy (Second Edition), Academic Press (2007).

27. O. H. S. Ollila, H. A. Heikkinen, H. Iwaï, Rotational Dynamics of Proteins from Spin Relaxation Times and Molecular Dynamics Simulations. J. Phys. Chem. B 122, 6559–6569 (2018).

28. R. Nencini, et al., Probing the dynamic landscape of peptides in molecular assemblies by synergized NMR experiments and MD simulations. Commun Chem 7, 28 (2024).

29. M. V. Fawaz, et al., Phospholipid Component Defines Pharmacokinetic and Pharmacodynamic Properties of Synthetic High-Density Lipoproteins. J Pharmacol Exp Ther 372, 193–204 (2020).

30. N. Sibille, P. Bernadó, Structural characterization of intrinsically disordered proteins by the combined use of NMR and SAXS. Biochemical Society Transactions 40, 955–962 (2012).

31. M. Chan-Yao-Chong, D. Durand, T. Ha-Duong, Molecular Dynamics Simulations Combined with Nuclear Magnetic Resonance and/or Small-Angle X-ray Scattering Data for Characterizing Intrinsically Disordered Protein Conformational Ensembles. J. Chem. Inf. Model. 59, 1743–1758 (2019).

32. T. Reddy, J. K. Rainey, Interpretation of biomolecular NMR spin relaxation parameters. Biochem. Cell Biol. 88, 131–142 (2010).

33. A. Debnath, L. V. Schäfer, Structure and Dynamics of Phospholipid Nanodiscs from All-Atom and Coarse-Grained Simulations. J. Phys. Chem. B 119, 6991–7002 (2015).

34. I. Siuda, D. P. Tieleman, Molecular Models of Nanodiscs. J. Chem. Theory Comput. 11, 4923–4932 (2015).

35. A. Y. Shih, I. G. Denisov, J. C. Phillips, S. G. Sligar, K. Schulten, Molecular Dynamics Simulations of Discoidal Bilayers Assembled from Truncated Human Lipoproteins. Biophysical Journal 88, 548–556 (2005).

36. L. R. Kjølbye, et al., General Protocol for Constructing Molecular Models of Nanodiscs. J. Chem. Inf. Model. 61, 2869–2883 (2021).

37. N. Sreerama, R. W. Woody, Estimation of Protein Secondary Structure from Circular Dichroism Spectra: Comparison of CONTIN, SELCON, and CDSSTR Methods with an Expanded Reference Set. Analytical Biochemistry 287, 252–260 (2000).

38. G. Zhu, Y. Xia, L. K. Nicholson, K. H. Sze, Protein Dynamics Measurements by TROSY-Based NMR Experiments. Journal of Magnetic Resonance 143, 423–426 (2000).

39. F. Delaglio, et al., NMRPipe: A multidimensional spectral processing system based on UNIX pipes. J Biomol NMR 6 (1995).

40. S. P. Skinner, et al., CcpNmr AnalysisAssign: a flexible platform for integrated NMR analysis. J Biomol NMR 66, 111–124 (2016).

41. P. Pernot, A. Round, R. Barrett, A. De Maria Antolinos, al., Upgraded ESRF BM29 beamline for SAXS on macromolecules in solution. J Synchrotron Rad 20, 660–664 (2013).

42. D. Orthaber, A. Bergmann, O. Glatter, SAXS experiments on absolute scale with Kratky systems using water as a secondary standard. J Appl Crystallogr 33, 218–225 (2000).

43. S. Förster, L. Apostol, W. Bras, Scatter : software for the analysis of nano- and mesoscale small-angle scattering. J Appl Crystallogr 43, 639–646 (2010).

44. K. Manalastas-Cantos, et al., ATSAS 3.0 : expanded functionality and new tools for small-angle scattering data analysis. J Appl Crystallogr 54, 343–355 (2021).

45. P. C. T. Souza, et al., Martini 3: a general purpose force field for coarse-grained molecular dynamics. Nat Methods 18, 382–388 (2021).

46. S. Jo, T. Kim, V. G. Iyer, W. Im, CHARMM‐GUI: A web‐based graphical user interface for CHARMM. J Comput Chem 29, 1859–1865 (2008).

47. K. Lindorff‐Larsen, et al., Improved side‐chain torsion potentials for the Amber ff99SB protein force field. Proteins 78, 1950–1958 (2010).

48. C. J. Dickson, et al., Lipid14: The Amber Lipid Force Field. J. Chem. Theory Comput. 10, 865–879 (2014).

49. J. Huang, et al., CHARMM36m: an improved force field for folded and intrinsically disordered proteins. Nat Methods 14, 71–73 (2017).

50. J. B. Klauda, et al., Update of the CHARMM All-Atom Additive Force Field for Lipids: Validation on Six Lipid Types. J. Phys. Chem. B 114, 7830–7843 (2010).

51. H. W. Horn, et al., Development of an improved four-site water model for biomolecular simulations: TIP4P-Ew. The Journal of Chemical Physics 120, 9665–9678 (2004).

52. H. J. C. Berendsen, D. Van Der Spoel, R. Van Drunen, GROMACS: A message-passing parallel molecular dynamics implementation. Computer Physics Communications 91, 43–56 (1995).

53. G. Bussi, D. Donadio, M. Parrinello, Canonical sampling through velocity rescaling. The Journal of Chemical Physics 126, 014101 (2007).

54. M. Parrinello, A. Rahman, Polymorphic transitions in single crystals: A new molecular dynamics method. Journal of Applied Physics 52, 7182–7190 (1981).

55. T. Darden, D. York, L. Pedersen, Particle mesh Ewald: An N ⋅log(N) method for Ewald sums in large systems. The Journal of Chemical Physics 98, 10089–10092 (1993).

56. C.-Y. Lu, D. A. Vanden Bout, Effect of finite trajectory length on the correlation function analysis of single molecule data. The Journal of Chemical Physics 125, 124701 (2006).

57. L. E. Kay, D. A. Torchia, A. Bax, Backbone dynamics of proteins as studied by nitrogen-15 inverse detected heteronuclear NMR spectroscopy: application to staphylococcal nuclease. Biochemistry 28, 8972–8979 (1989).

58. K. T. Dayie, G. Wagner, J.-F. Lefèvre, Theory and Practice of Nuclear Spin Relaxation in Proteins. Annu. Rev. Phys. Chem. 47, 243–282 (1996).

59. D. Svergun, C. Barberato, M. H. J. Koch, CRYSOL – a Program to Evaluate X-ray Solution Scattering of Biological Macromolecules from Atomic Coordinates. J Appl Crystallogr 28, 768–773 (1995).

60. D. I. Svergun, et al., Protein hydration in solution: Experimental observation by x-ray and neutron scattering. Proc. Natl. Acad. Sci. U.S.A. 95, 2267–2272 (1998).

61. W. Kabsch, C. Sander, Dictionary of protein secondary structure: Pattern recognition of hydrogen‐bonded and geometrical features. Biopolymers 22, 2577–2637 (1983).

62. R. Gowers, et al., MDAnalysis: A Python Package for the Rapid Analysis of Molecular Dynamics Simulations in (2016), pp. 98–105.

63. J. García De La Torre, M. L. Huertas, B. Carrasco, Calculation of Hydrodynamic Properties of Globular Proteins from Their Atomic-Level Structure. Biophysical Journal 78, 719–730 (2000).

