## Supplementary information for "Exploring Peptide Nanodiscs Structure and Dynamics through Synergistic Approach of NMR Spectroscopy, SAS and MD Simulations"

[1]Division of Pharmaceutical Biosciences, Faculty of Pharmacy, University of Helsinki, Helsinki, Finland
[2]Institute of Biotechnology, University of Helsinki, Helsinki, Finland
[3]VTT Technical Research Centre of Finland, Espoo, Finland
*Corresponding author

**This PDF file includes:**

Figures S1 to S5
Tables S1 to S2

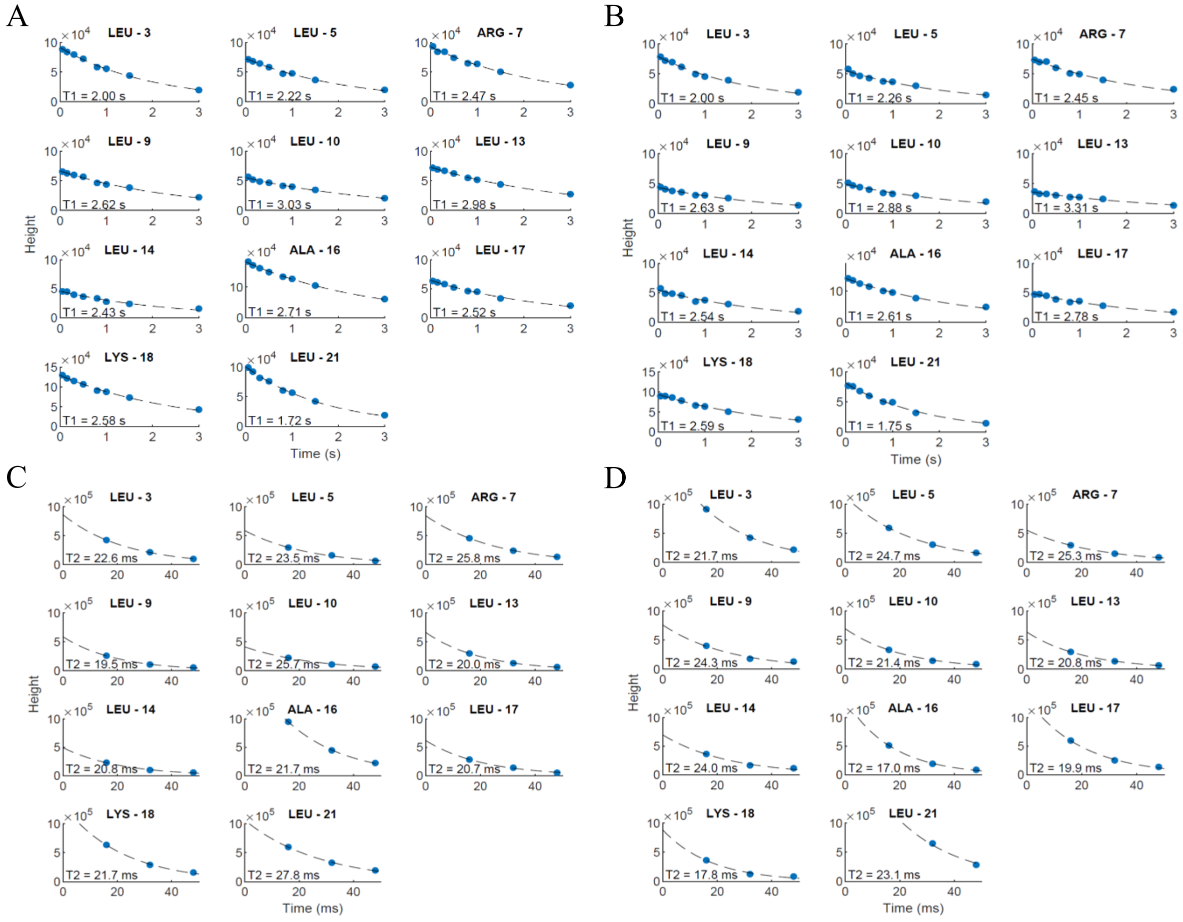


**Figure S1. NMR relaxation parameter decays for the duplicate samples.** (A) $T_1$ decays for sample 1. (B) $T_1$ decays for sample 2. (C) $T_2$ decays for sample 1. (D) $T_2$ decays for sample 2.

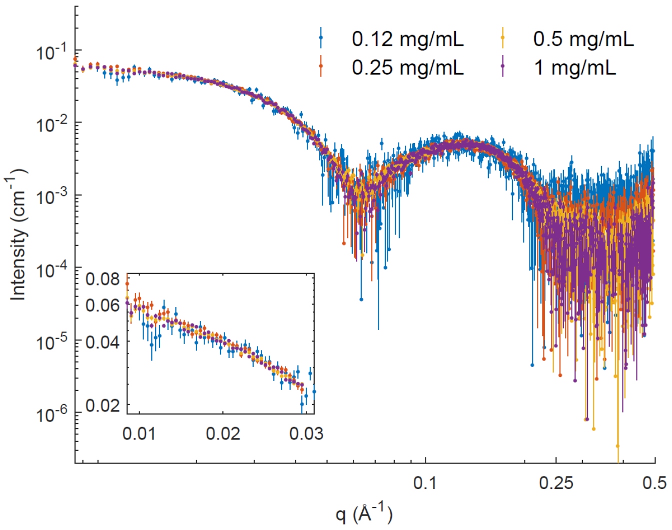


**Figure S2. Influence of the peptide concentration in nanodiscs on the SAXS intensities.**

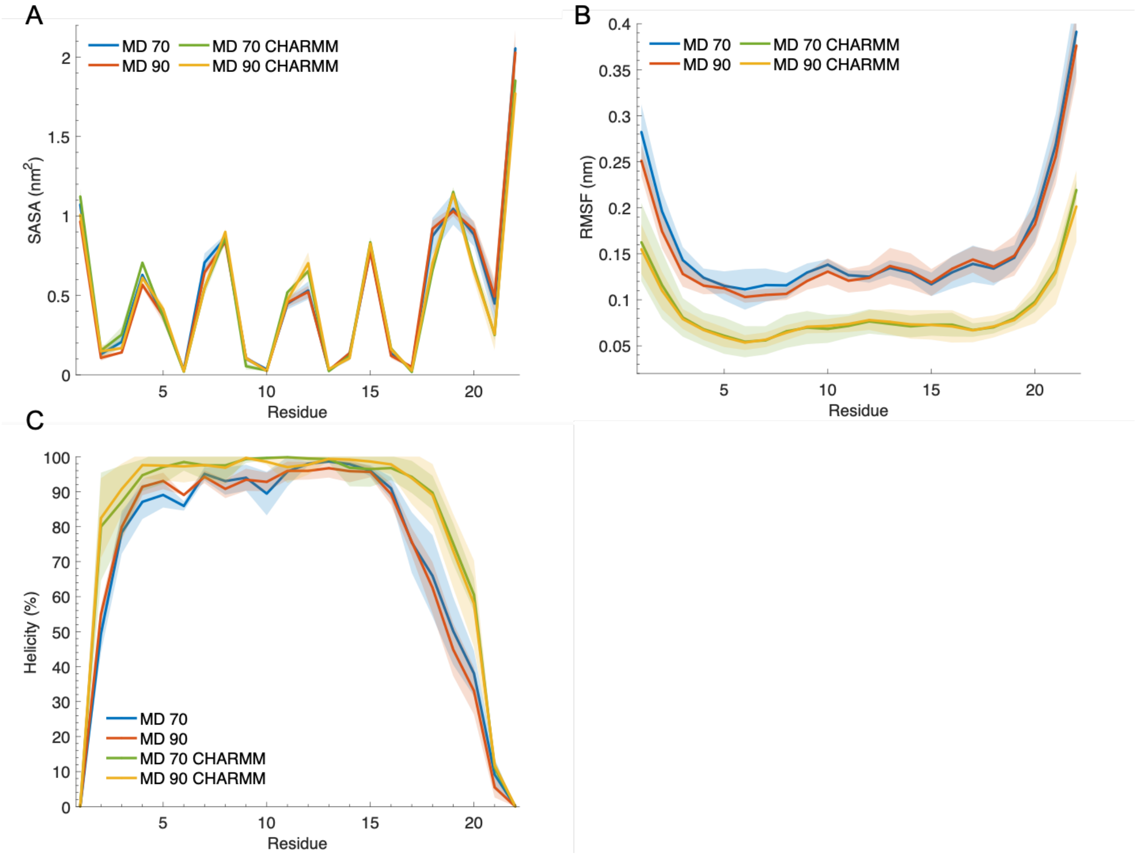


**Figure S3. Peptide behavior under different force fields.** (A) Solvent accessible surface area (SASA) per residue. (B) Root mean square fluctuation (RMSF) per residue. (C) Helicity per residue. MD data are represented as mean ± SD of triplicate simulations.

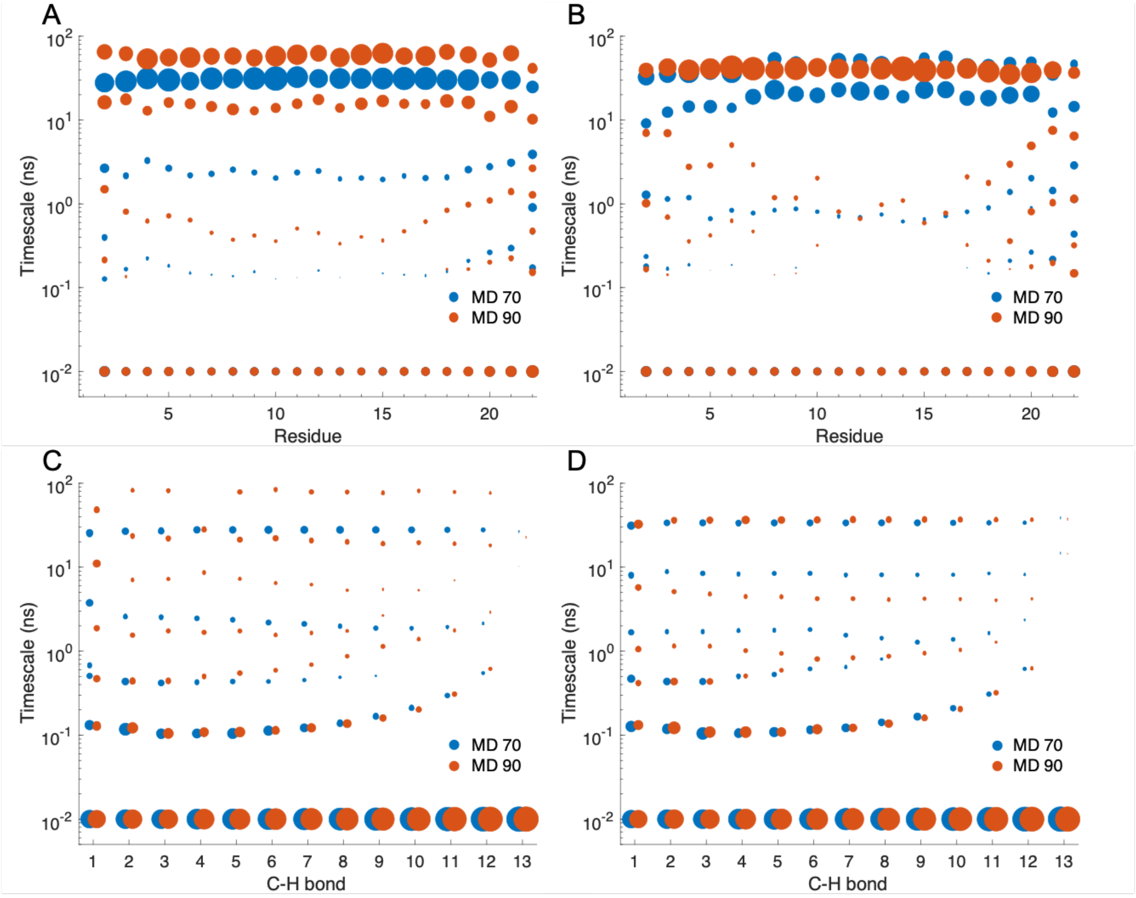


**Figure S4. Dynamic landscape of 22A peptides and DMPC phospholipids.** Dynamic landscapes of 22A peptides for the second replicate (A) and the third replicate (B). Dynamic landscapes of DMPC phospholipids for the second replicate (C) and the third replicate (D). C-H bond 13 refers to the terminal methyl group of DMPC phospholipids. The point sizes represent the weight of each timescale in the rotational relaxation process.

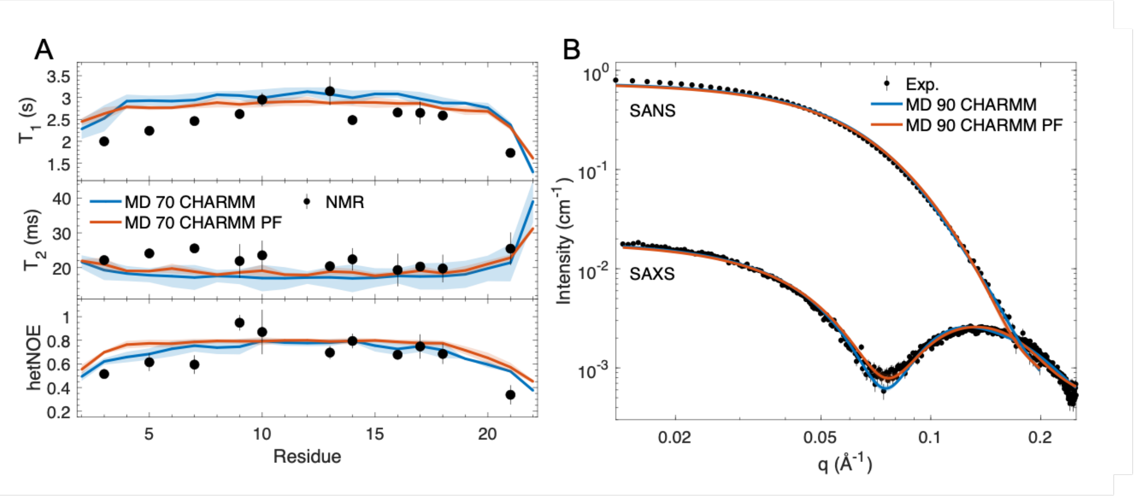


**Figure S5. Effect of the orientation of peptides in the starting structure of nanodisc on the reproduction of experimental data from MD simulations.** (A) NMR relaxation parameters. (B) SAXS and SANS curves. MD data are represented as mean ± SD of triplicate simulations.

| Force field | Starting configuration | System size (number of lipids) | Box size in X, Y and Z dimensions | Length of rotacf analyzed | Simulated time and number of replicates |
|---|---|---|---|---|---|
| AMBER | MARTINI final snapshot | 30 | 10 nm | 50 ns | 3 μs x 1 |
| | | 50 | 11 nm | 50 ns | 3 μs x 3 |
| | | 70 | 12 nm | 100 ns | 6 μs x 3 |
| | | 90 | 14 nm | 100 ns | 3 μs x 3 |
| | | 110 | 16 nm | 100 ns* | 3 μs + 1 μs x 2 |
| | Parallel picket fence | 70 | 12 nm | - | 2 μs x 3 |
| | | 90 | 14 nm | - | 1 μs x 3 |
| CHARMM | MARTINI final snapshot | 70 | 12 nm | 100 ns | 3 μs x 3 |
| | | 90 | 14 nm | - | 1 μs x 3 |
| | Parallel picket fence | 70 | 12 nm | 100 ns | 3 μs x 3 |
| | | 90 | 14 nm | - | 1 μs x 3 |

*only the 3 μs trajectory was included in analysis

**Table S1. Performed MD simulations.**

| | MD CHARMM | MD CHARMM PF |
|---|---|---|
| $\chi^2$ - T1 | 0.68 | 0.59 |
| $\chi^2$ - T2 | 0.68 | 0.58 |
| $\chi^2$ - hetNOE | 0.74 | 0.93 |
| $\chi^2$ - SAXS | 4.10 | 8.46 |
| $\chi^2$ - SANS | 5.50 | 8.35 |

**Table S2. Agreement between MD simulations and experiments for the CHARMM force field.**
